# Neurophysiological excitation/inhibition imbalance in young adults burdened with childhood interpersonal trauma

**DOI:** 10.64898/2026.01.14.699432

**Authors:** Alejandro Orozco Valero, Natalia Kopiś-Posiej, Víctor Rodríguez-González, Víctor Gutiérrez-de Pablo, Christian Morillas, Jesús Poza, Carlos Gómez, Pablo Martínez-Cañada, Paweł Krukow

**Author notes:** Corresponding authors. These authors jointly supervised this work. **Email:**. **Author Contributions:** A.O.V., N.K.P., V.R.G., J.P., C.G., P.M.C., and P.K. conceptualized the study and developed the methodology; A.O.V., N.K.P., P.M.C., and P.K. conducted the investigation; all authors performed formal analysis; A.O.V., P.M.C., and P.K. wrote the original draft; all authors reviewed and edited the manuscript. **Competing Interest Statement:** The authors declare no competing interest.

## Abstract

Adverse childhood experiences, such as violence, abuse, and neglect, are increasingly recognized as significant modifiers of brain development. Here, we tested whether adults with histories of childhood trauma, but without psychiatric comorbidities exhibit altered excitation/inhibition (E/I) balance, as indicated by electroencephalography (EEG) signatures. Participants, divided into low-trauma and high-trauma groups, underwent two experimental conditions: eyes-closed resting-state recording and a reaction-time task with visual stimuli. From these data, we computed 1/f spectral slopes, a widely used electrophysiological marker of E/I balance; we complemented these analyses with a leaky integrate-and-fire (LIF) microcircuit model combined with a biophysically grounded forward-modeling approach to simulate realistic brain signals and derive E/I balance estimates. Group comparisons for both slopes and E/I estimates revealed significant resting-state differences, characterized by a shift toward increased neuronal excitation in the high-trauma group. The high-trauma group exhibited altered stimulus-related 1/f slope dynamics relative to the pre-stimulus baseline, reflecting attenuated neuronal inhibition. E/I ratio measures were not significantly correlated with participants’ transient affective states. Together, these findings suggest that childhood trauma is associated with enduring, trait-like alterations in cortical E/I balance that extend beyond affective state and manifest across both resting and task-related brain dynamics.

**Significance Statement:** Childhood trauma is a major risk factor for mental illness, yet its lasting effects on basic brain physiology remain poorly understood. Using electroencephalography combined with biophysically grounded neural circuit modeling, we show that young adults with a history of childhood interpersonal trauma exhibit a persistent shift toward cortical hyperexcitation at rest and a reduced ability to engage inhibitory control during cognitive processing. These effects occur even in individuals without psychiatric diagnoses and are independent of current anxiety or depression, indicating a trait-like neurophysiological footprint of early adversity. This excitation-inhibition imbalance may represent a transdiagnostic vulnerability linking childhood trauma to later psychopathology.

## Introduction

The potential consequences of psychologically negative childhood experiences on adult functioning and brain health are the subject of research in the fields of psychiatry, epidemiology, and neuroscience (1–3). It is increasingly recognized that bearing violence, abuse, and neglect at a younger age significantly modifies the developmental trajectories of the nervous system (4), biological mechanisms of the stress response (5), emotion regulation (6), and cognitive processes (7, 8). There is accumulating evidence suggesting that this influence may be long-lasting and constitute a significant risk factor for the emergence of mental disorders, accelerated cognitive decline (9), and poorer somatic health in adulthood (10–12).

A growing body of neuroimaging evidence suggests that exposure to early-life adversity is associated with complex and widespread alterations in brain structure and function that persist in adulthood (13, 14). However, the inclusion of individuals with formal psychiatric diagnoses in study samples limits the ability to unequivocally attribute these neural alterations specifically to childhood trauma. Well-controlled resting-state and task-based functional Magnetic Resonance Imaging (fMRI) studies indicate that adults burdened by childhood trauma (ChT) exhibit limbic system hyperactivity, including amygdala hyperexcitability, alongside reduced frontal activation and insufficient prefrontal engagement during working memory tasks (15, 16). In contrast, other neuroimaging findings report the opposite pattern, showing increased recruitment of metabolic brain resources in ChT adults when coping with tasks of varying difficulty compared with non-exposed individuals (17). Collectively, these findings point to substantial heterogeneity within the ChT group; rather than a single recurring pattern, the evidence suggests dysregulated or maladaptive neuronal and metabolic responses to task demands and resting-state conditions, reflecting difficulties in maintaining neural homeostasis (18, 19).

Although initially appearing unrelated, some neurobiological alterations noticed in the ChT group might have a common background. A rearranged proportion of brain activity frequencies related to the state of relaxation and vigilance (20, 21), an abnormally increased orientation reflex during basic information processing (22), and signs of diminished prefrontal cortex top-down control over subcortical structures (23, 24), collectively suggest that ChT-burdened adults might present with an imbalance between neuronal excitation (E) and inhibition (I). The balance between these processes is considered a major factor determining healthy neuronal development, especially during critical periods of experience-dependent plasticity (25). Animal studies, e.g., (26), show that the excitation/inhibition (E/I) ratio of sensory neurons becomes optimally tuned by environmental stimulation. Alterations in E/I balance are well-documented in neuropsychiatric disorders associated with early-onset psychopathology, such as autism (27–29). E/I balance is refined by the development of GABAergic parvalbumin-positive (PV) interneurons, providing a cellular structure of inhibitory circuitry reducing stimulus-irrelevant activation enhancing signal-to-noise ratio (30–32). Human fMRI neuroimaging studies indicate that E/I balance matures towards effective inhibition, which reduces neuronal noise generated by uncontrolled overactivity (33). Such modelling of the E/I ratio towards effective inhibition is already proven determinant of proper neurocognitive development (25).

One of the clues linking ChT and E/I imbalance is a series of epidemiological observations (34) and animal model studies indicating a relationship between adverse childhood experiences (ACE) and epilepsy, or generally ictal features of the neurophysiological activity. Research utilizing rodent models revealed that experimental settings, such as early maternal deprivation intensify status epilepticus, lead to the amplification of the N-methyl-D-aspartate (NMDA) in the postnatal phase and the amygdala kindling mechanism (35–37). Additionally, Dube et al. (38) in a group of Spraguee Dawley rats subjected to chronic early-life stress observed an increased proportion of spike series in electroencephalography (EEG) records and a significant increase of amygdalar corticotropin-releasing hormone (CRH) considered as pro-convulsant neuropeptide produced in response to stress. A series of studies also indicate that young rats exposed to various forms of stress develop more severe seizure phenotypes following administration of pro-epileptic agents and are more likely to develop chronic epilepsy, an outcome not observed in non-stressed rats despite comparable pro-convulsant exposure (39, 40). It has also been suggested that, in humans, there are enduring relationships between stress-induced remodeling of the hypothalamic-pituitary-adrenal (HPA) axis and the risk of epilepsy (41); moreover, early stress disrupts neurogenesis in selected areas of the hippocampus, which plays an important role in inhibiting the HPA axis (42). Subsequent atrophic changes in the hippocampus can increase the expression of CRH mRNA leading to an increase in circulating corticosterone, a pro-seizure substance (43, 44). Ultimately, Karst et al. (45) showed that early life stress alters the developmental phases of the basic neurochemical processes responsible for the neuronal E/I balance in mice.

Finally, several preliminary studies used Proton Magnetic Resonance Spectroscopy (^1^HMRS) in ChT to assess a profile of neurochemical markers accounting for neuronal excitation and inhibition processes. For example, Averill et al. (46) found a significant and positive correlation between early life adversity and glutamine levels in adults with major depressive disorder. Sonmez et al. (47), in a sample of adolescents and young adults, obtained results suggesting increased excitability, indexed by NAA/Glx ratio downturn within the anterior cingulate cortex. Further correlational analyses showed that a significant association between ChT and spectroscopy outcomes, primarily within the subsample of participants with depression. Relationships between adverse childhood experiences and alterations regarding the GABAergic system were corroborated by at least two independent studies (48, 49), although it should be noted that the found associations were mainly correlational, and some groups involved participants with confirmed affective disorders. MRS is one of the few imaging techniques that enables in vivo assessment of neurochemical indices; however, it also has several limitations, i.e., the inability to evaluate the dynamics of these indices and the restriction of measurements to relatively narrow neuronal structures. The concentration of GABA on the MRS spectra is low compared with larger peaks, such as those associated to creatine (Cr) and choline (Cho). Furthermore, the locations of glutamate and GABA on the spectrum are close, so the exact measurement might be compromised, especially when 1.5 Tesla MRI equipment is utilized (50). In mentioned studies, MRS measurement was limited to a single structure or at least a few preselected brain areas.

Given the challenges inherent in measuring E/I imbalance in vivo, indirect methods are required. EEG and magnetoencephalography (MEG) signals enable E/I balance mapping over the entire cortical surface and, compared with MRS, track E/I changes with higher temporal resolution (51). Importantly, these electrophysiological signals allow for the characterization of scale-free neural activity, from which indices of E/I balance, such as the aperiodic exponent of the 1/f spectral power law (“1/f slope”), can be derived. Although informative, such EEG/MEG-derived markers remain phenomenological and do not uniquely specify the underlying synaptic mechanisms. In this context, the combination of machine-learning algorithms with brain model simulations becomes useful, as it enables the mapping of observed neural dynamics onto candidate cortical circuit parameters (52, 53). These computational frameworks, informed by EEG- and MEG-derived neurophysiological data among other complementary modalities, facilitate mechanistic interpretation of E/I-related spectral features. Notably, the 1/f slope has been validated across multiple clinical populations and in relation to developmental changes in E/I balance (27, 28, 30, 54–56); however, to our knowledge, it has not yet been applied to investigate E/I balance in adults burdened with ChT. Considering that ChT is an important risk factor for many adverse health conditions (10, 12) and the fact that previous ChT-related studies on E/I balance are sparse and based on small-numbered or highly heterogeneous groups, our study aimed to examine the major assumption that ChT leads to a relatively persistent imbalance in excitatory and inhibitory neuronal mechanisms noticeable in adults burdened with such experiences.

Specifically, we studied whether there are significant differences in E/I indicators between groups contrasted by the degree of ChT exposure. These analyses considered both eyes-closed resting-state and task-related conditions. The study by Gyurkovics et al. (57) showed that stimulus presentation induces a shift in the E/I balance toward inhibition. Therefore, the analysis of the results also involved assessing the interaction effect regarding the group and the reconfiguration of the E/I ratio pattern in response to the stimuli. Considering previous empirical literature, our major hypothesis is that the ChT group will exhibit E/I shift towards increased neuronal excitation and/or diminished inhibition. To the best of our knowledge, this is the first time that these analyses are performed in ChT adults so far; therefore, no available data point out specific cortical regions in which these changes should be observed. However, given previous findings on early adversities-related prefrontal cortex top-down control disruptions (23, 24), we hypothesize that task-related E/I alterations in the ChT group will occur primarily in the anterior cortical regions. The involvement of participants with psychiatric disorders in some previous studies examining the relationship between ChT and brain function may have hindered the ability to definitively isolate the effects of ChT on specific neurophysiological activity (48, 49). Therefore, the present study included only individuals from the general population, with no diagnosed mental disorders or history of psychopharmacological treatment.

Finally, ChT-exposed adults, even in the absence of clinically severe affective disturbances, often report elevated levels of anxiety and depressive symptoms (58, 59). Moreover, experimental induction of negative affect has been shown to shift the E/I balance toward neuronal hyperexcitation (60), supporting the need to control for affective status in the currents study. Given the considerable body of research evidence demonstrating the persistence of the neural effects of early adversities (15, 61), we hypothesize that trauma-related alterations in E/I balance represent a trait-like neural characteristic that is dissociable from state-dependent fluctuations in affect. Accordingly, we expect no statistically significant correlations between E/I balance measures and levels of depressiveness or anxiety within the ChT-exposed sample.

## Materials and Methods

### Overview of our model-based inference approach

Our goal was to analyze EEG data from participants with different levels of ChT and infer possible imbalances in E/I ratio. To achieve this, we developed a model-based inference framework that links changes in EEG signals to underlying shifts in E/I. Fig. 1 provides an overview of the full methodological pipeline, which we summarize here before describing each step in detail in the following sections.

**Figure 1.**
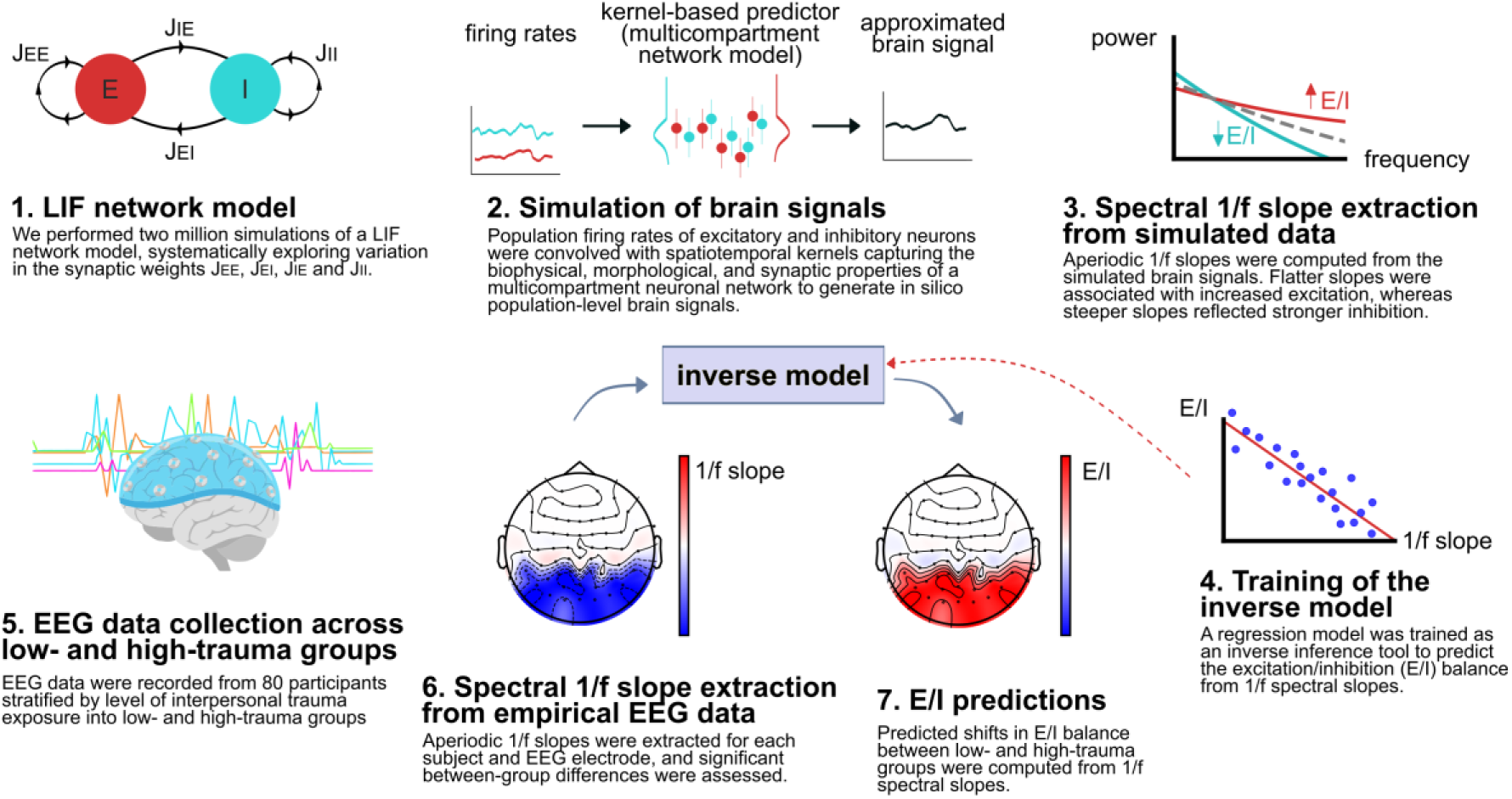
Modelling and inference pipeline. Schematic illustration of the workflow, from cortical circuit simulations to regression-based inference on real data. See the main text for a detailed explanation of each step.

We began by simulating a local cortical circuit model based on leaky integrate-and-fire (LIF) neurons with recurrent connections between excitatory and inhibitory populations (step 1). From the resulting spiking activity, we applied a state-of-the-art biophysically grounded kernel method to compute the population-level brain signal (62) (step 2). This kernel was derived using a hybrid modeling approach that incorporates an equivalent multicompartment network model, allowing us to account for biophysical, morphological and syntactic properties of the network such as synapse locations and cell distributions. From simulated signals, we extracted the 1/f spectral slope (step 3), a feature that has been consistently associated with E/I balance in the literature: flatter slopes indicate increased excitation, while steeper slopes reflect shifts toward inhibition. Then, we trained a Ridge regression model on the simulated data to construct an inverse mapping from EEG features to underlying E/I changes (step 4). Finally, we applied this model to our empirical EEG recordings from two groups differing in the ChT exposure (step 5), using the extracted spectral slopes (step 6) as input and obtaining predictions of E/I alterations potentially linked to ChT (step 7).

### EEG dataset of childhood trauma

The final dataset includes 80 participants sampled concerning the level of interpersonal trauma exposure, but without a current diagnosis of post-traumatic stress disorder (PTSD), according to the score obtained in the Impact Event Scale-Revised (63). The study participants came from the general population, aged 19-25 yrs. They were selected from a larger group of volunteers, mainly students. Volunteers could take part in the main study if they received a score within a predetermined Childhood Trauma Questionnaire total points range (CTQ (64)), and if none of the following exclusion criteria were found: previously diagnosed mental disorders, history of psychopharmacological treatment, neurological diseases, including neuroinfections, traumatic injuries, and epilepsy. Based on participants’ CTQ total scores and established cut-off values distinguishing low and moderate-to-severe trauma exposure, participants were stratified into two groups: a low-trauma group (n = 41, CTQ total score ≤ 36) and a high-trauma group (n = 39, CTQ total score ≥ 51). The resulting groups did not differ significantly with respect to basic demographic characteristics: age (22.70 yrs, SD = 1.86) in the low-trauma group versus 22.38 yrs (SD = 2.21) in the high-trauma group; proportion of males (23.07% vs. 34.14%); and years of education (14.89, SD = 2.44 vs. 14.66, SD = 2.00, respectively; all *p* ≥ 0.10).Additionally, all participants fulfilled Beck Depression Inventory (BDI-II (65)), and State-Trait Anxiety Inventory (STAI (66)) to evaluate their affective status. The application of these questionnaires served to exclude participants with scores indicating clinically relevant depression and anxiety disorders. Informed consent was obtained from all individual participants; the study followed the Declaration of Helsinki and was approved by the local ethical committee (consent KE-0254/202/ 09/2023).

EEG data were recorded using a 64-channel Ag/AgCl actiCap active electrodes (Brain Products, GmbH, Germany), arranged according to the international 10-10 system, under two different conditions. The first condition included 5-minute resting-state eyes-closed recordings. The second condition was a visual reaction-time task lasting approximately 30 minutes. The main task was to react to the target stimuli, which were rectangles with vertical or horizontal stripes displayed in the center of the screen. Depending on whether the stripes’ orientation was horizontal or vertical, participants had to press the right or left key, respectively, as fast and as accurately as possible. The procedure began with a short training session during which ten target stimuli were presented, and participants received feedback (’good/bad’/’faster’) regarding the correctness of their response. The experiment was designed in two randomized blocks differing in the time interval between the appearance of the fixation cross and the display of the target stimulus. In one block, a fixation cross was displayed for 2000-2500 ms (short interval block), while in another, it was 7000-9000 ms (long interval block). The trials began with the presentation of a fixation dot (750 ms, centered on the screen), followed by a fixation cross (either a short or long version), and then a target stimulus. The target was displayed until response, or for a maximum of 2000 ms. In total, 250 cue-target trials were used in the entire procedure. To capture the evolution of the neural response to the stimulus, from early sensory processing to later cognitive stages, we employed two types of 500 ms epochs to analyze each trial: a 500 ms pre-stimulus baseline and a 500 ms post-stimulus period. Task design with two main blocks was intended to reduce monotony and, on the other hand, introduce an experimental factor that generates attentional lapses (67) expected specifically in ChT individuals, given the cognitive deficits documented in this group (68). Performance metrics, including mean reaction time, its variability, and responses accuracy, were evaluated for each task block.

### Computational modelling of cortical circuit dynamics and brain signals

To simulate cortical circuit dynamics, we used a spiking neural network model consisting of a population of 8192 excitatory (E) and 1024 inhibitory (I) leaky integrate-and-fire neurons, driven by an external fixed-rate Poisson input (62). We defined a 12000 ms simulation configuration with a 0.0625 ms time step, discarding the first 2000 ms as transient time in the analysis. We performed two million simulations using this model, systematically varying the synaptic weights *J_EE_*, *J_EI_*, *J_IE_*, and *J_II_*, where *J_YX_* denotes the synaptic strength from presynaptic population *X* to postsynaptic population *Y*. The remaining variables of the network were fixed using the best-fit parameters employed in the corresponding publication (62). We employed the kernel computation method (62) to calculate the population extracellular signal, which was used as a representation of EEG activity. Accordingly, spike rates were convolved with spatiotemporal filter kernels that incorporate neuronal biophysics, the spatial and temporal distribution of cells and synapses, and the connectivity of network-equivalent populations of conductance-based multicompartment neurons. We used the same network of ball-and-stick neurons, with the same network configuration, as in (62).

### Spectral slopes derived from EEG data

We parameterized the power spectrum of each EEG signal (both simulated and empirical) to extract its spectral features. This was accomplished using *specparam* (formerly *fooof*; (69)), a widely used library that models neural power spectra by separating them into two distinct components: periodic oscillations (which appear as peaks over the background) and the aperiodic 1/f-like component. Our primary feature of interest for this analysis was the exponent of the aperiodic component, commonly referred to as the spectral slope. The *specparam* model was configured to fit the power spectrum across a frequency range of 5 to 45 Hz, a frequently used frequency range according to recent guidelines (70). To ensure the reliability of the derived parameters, we implemented a strict quality control step: any model fit with an R-squared (R^2^) value below 0.9 was excluded from further analysis. Other parameters of the spectral fitting were: ’peak_threshold’: 1, ’min_peak_height’: 0, ’max_n_peaks’: 5, ’peak_width_limits’: (10, 50) Hz.

### Inference of cortical circuit parameters

A linear regression model with Ridge regularization based on the *ncpi* library (52) was trained on the 2 million simulated samples to infer circuit E/I from EEG-derived features. This approach, referred to as inverse modeling (52), learns the mapping between observable EEG features and the underlying cortical circuit parameters. Once validated on the simulation dataset where the ground truth is known, this trained model serves as an inference tool that can be applied to empirical EEG data to estimate the most probable circuit parameters that gave rise to the observed signals. To enhance generalization, ensure robustness to random seed selection and optimize regularization strength, we performed 20 repetitions of a grid-search-based 10-fold cross-validation, resulting in a total of 200 folds. After the training process was finished, the E/I ratio was then computed from inferred synaptic weights *J_EE_*, *J_EI_*, *J_IE_*, and *J_II_* as follows:

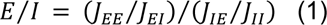

This summary metric quantifies the net balance between excitatory and inhibitory processes in the circuit model and can be interpreted as a global E/I ratio derived from simulated data.

### Statistics

Group differences in demographic characteristics and questionnaire outcomes were analyzed using parametric *t*-tests or χ² tests for categorical variables. Cognitive task outcomes were coded as the level of responses accuracy and selected features of the reaction time (RT) distribution. RT data were analyzed using both traditional parametric metrics (mean and within-subject standard deviation) and ex-Gaussian parameters. The ex-Gaussian model represents the joint contribution of normal and exponential components through three parameters: *µ* (mu), corresponding to the mean of the Gaussian component; *σ* (sigma), a standard-deviation–like index reflecting the variability of shorter and longer RTs around the mean; and *τ* (tau), representing the mean and variability of the exponential component. Parameter estimation was conducted on the pre-cleaned RT data in accordance with the guidelines of Lacouture and Cousineau (71), using the MATLAB toolbox DISTRIB (version R2017a; MathWorks Inc., Natick, MA, USA).

The Anderson-Darling test was employed to assess whether the distributions of extracted slope features and inferred E/I ratios conformed to normality for each experiment. As a complementary assessment, Q-Q plots were generated for each EEG electrode to visually inspect the distributional assumptions. For samples meeting the normality criterion, parametric comparisons were conducted using the Student’s *t*-test. To quantify the magnitude of group differences, effect sizes were calculated using Cohen’s *d*. We report only statistically significant results (*p* < 0.05) with a Cohen’s *d* greater than 0.2.

Additionally, correlations between inferred E/I ratios and questionnaire scores were computed using Spearman’s rank correlation coefficient (*r*). To assess statistical significance, a permutation test was applied to estimate the associated *p*-values. We report only statistically significant results (*p* < 0.05).

## Results

### Behavioral outcomes

As expected, the high-trauma group scored significantly higher on the CTQ compared with low-trauma group (58.66 *vs.* 34.85, *p* < 0.0001; Table 1). According to CTQ cut-off thresholds (64), all participants included in the low-trauma group can be assigned as having non-to-low trauma exposure, while all participants from the high-trauma group were burdened with moderate-to-severe exposure. Compared with the low-trauma group, the high-trauma group had significantly higher RT variability as indexed by the sigma parameter in both task blocks and significantly lower response accuracy in the block with a shorter pre-stimulus interval (Table 1).

**Table 1.** Behavioral data on the study groups.

| ***Behavioral data*** | Low-trauma<br>(*n* = 41) |  | High-trauma<br>(*n* = 39) |  | *t/χ<sup>2</sup>* | *p* |
| --- | --- | --- | --- | --- | --- | --- |
|  | *M* | *SD* | *M* | *SD* |  |  |
| *Questionnaire data* |  |  |  |  |  |  |
| CTQ total | 34.85 | 2.27 | 58.66 | 8.03 | -13.979 | < **0.0001** |
| STAI state | 28.21 | 8.46 | 33.64 | 8.85 | -2.799 | **0.006** |
| STAI trait | 29.24 | 9.98 | 37.28 | 9.53 | -3.679 | **0.0004** |
| BDI-II total | 7.73 | 8.70 | 16.02 | 9.51 | -4.070 | **0.0001** |
| *Task-related data: short interval block* |  |  |  |  |  |  |
| mean RT | 593.69 | 89.33 | 607.05 | 90.60 | -0.664 | 0.508 |
| *μ* | 480.41 | 59.78 | 479.47 | 63.14 | 0.068 | 0.945 |
| *σ* | 52.99 | 15.18 | 62.12 | 19.80 | -2.320 | **0.022** |
| *τ* | 113.76 | 45.06 | 128.07 | 47.40 | -1.384 | 0.170 |
| Accuracy | 0.98 | 0.01 | 0.97 | 0.01 | 2.115 | **0.037** |
| *Task-related data: long interval block* |  |  |  |  |  |  |
| mean RT | 653.13 | 110.74 | 672.58 | 112.20 | -0.780 | 0.437 |
| *μ* | 522.44 | 72.43 | 540.59 | 81.72 | -1.052 | 0.295 |
| *σ* | 56.21 | 19.74 | 67.03 | 25.26 | -2.140 | **0.035** |
| *τ* | 132.48 | 58.51 | 134.41 | 54.17 | -0.153 | 0.878 |
| Accuracy | 0.74 | 0.01 | 0.73 | 0.01 | 0.747 | 0.457 |

### Predictions of E/I imbalances

The first step of the analysis was to compare the low- and high-trauma groups in terms of differences in resting-state EEG measures (Fig. 2A). Group comparisons revealed a significant decrease in the absolute 1/f slope values (|*d*| > 0.2, *p* < 0.05) in the high-trauma group compared to the low-trauma group. This effect, which indicates a flattening of the power spectrum slope, was localized over posterior electrodes. Next, an inverse model, derived from simulated data, was employed to infer the E/I balance from the empirical EEG signals (see *Material and Methods* for details). Consistent with the spectral slope analysis, the model predicted a significant increase in the E/I ratio within the same posterior regions for the high-trauma group.

**Figure 2.**
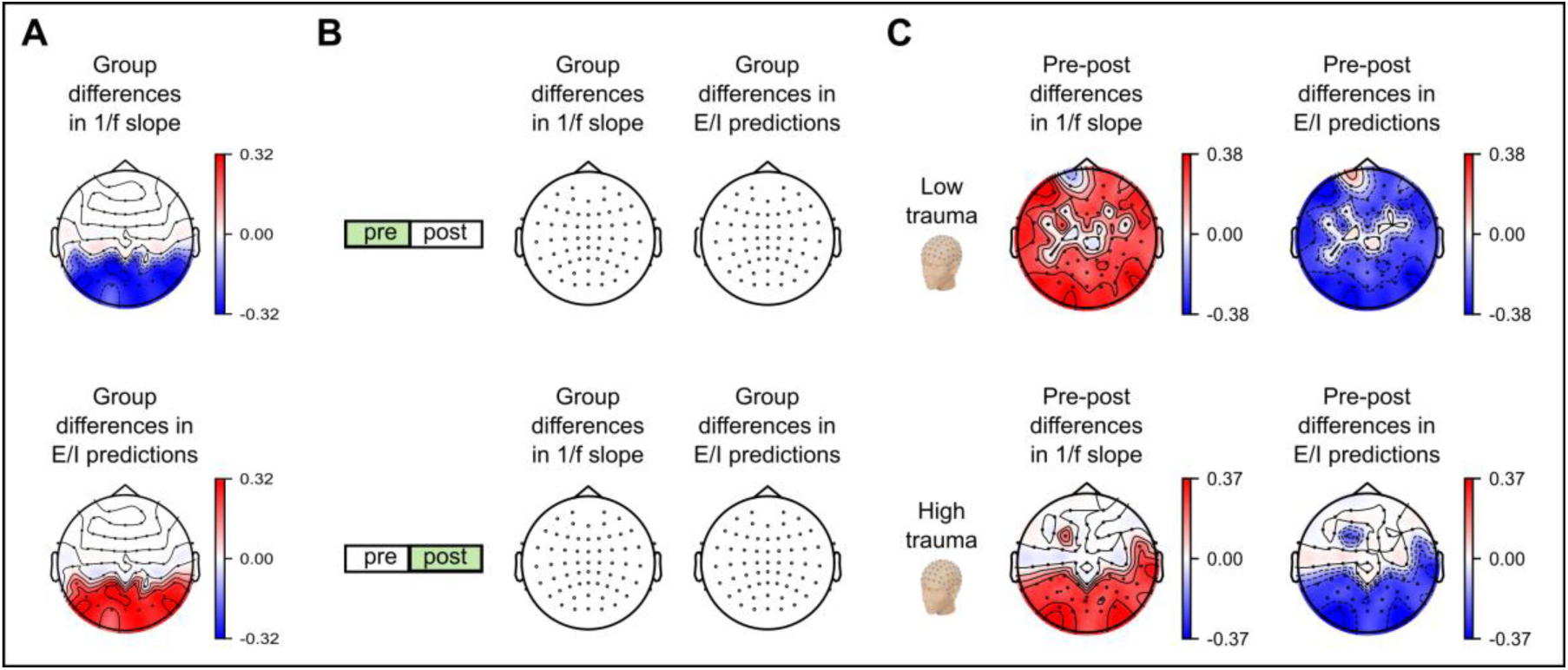
Resting-state and task-related regional differences in 1/f slope and inferred E/I balance. Brain topographies illustrating group differences between low- and high-trauma groups, expressed as Cohen’s *d*, for 1/f spectral slopes and model-based E/I estimates. Panel A shows resting-state results, while panel B shows task-related results (calculated from pre- and post-stimulus periods). Panel C displays differences (Cohen’s d) between pre- and post-stimulus measures within each group. Effect sizes are displayed only for regions in which |*d*| > 0.2 and differences reached statistical significance (*p* < 0.05) for both 1/f spectral slopes and model-based E/I predictions. Results are calculated using the short interval block.

For the task-related data (Fig. 2B and C), the analysis was split into pre- and post-stimulus segments to enable two types of comparisons: (i) differences between the low- and high-trauma groups (inter-group); and (ii) changes within each group from pre- to post-stimulus (intra-group). The direct comparison between groups revealed few significant differences (Fig. 2B and Supp. Fig. 1). In both the pre- and post-stimulus segments, only a small number of electrodes showed substantial group differences in either the spectral slope or the predicted E/I balance, and these were observed only in the long-interval block (Supp. Fig. 1). In contrast, analyses of stimulus-driven changes (pre- vs. post-stimulus, Fig. 2 C) revealed prominent effects. A significant widespread steepening of the spectral slope (i.e., an increase in its absolute value) was observed across the cortex in the low-trauma group, mirrored by a similar broad shift in the E/I balance toward inhibition. Compared to the high-trauma group, a differential group-specific pattern emerged: frontal electrodes in the low-trauma group showed strong stimulus-related modulation, whereas such an E/I ratio modulation was absent in the high-trauma group.

### Correlations with psychological variables

The next analysis involved computing the correlations between the mean E/I ratio predictions and the scores from questionnaires assessing affective characteristics of the study subjects. Similar correlation patterns are expected for the spectral slope given the linear relationship between slopes and inferred E/I values. Figure 3 shows results of this analysis performed separately for low-trauma group, high-trauma group, and for all participants combined in one sample. Correlations were computed using posterior electrodes, corresponding to regions that showed significant group differences in 1/f slope and inferred E/I ratio. None of the analysed correlation measures reached the statistical significance threshold of *p* < 0.05.

**Figure 3.**
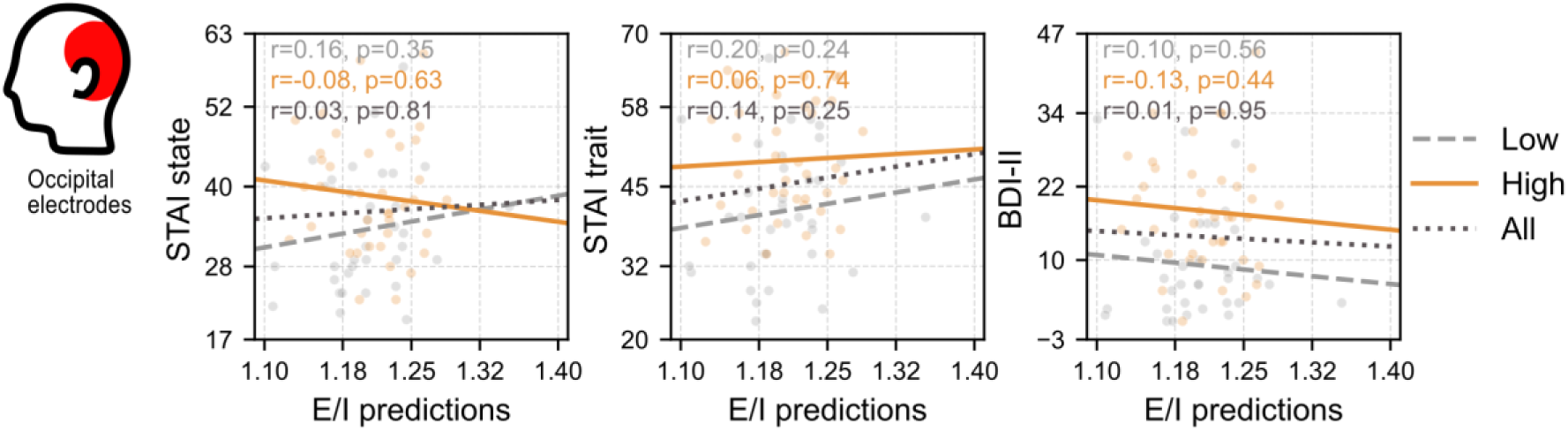
Correlations between inferred resting-state E/I values and selected psychological measures. Spearman’s correlations (*r*) and statistical significance levels (*p*) encompassing E/I predictions and psychological questionnaire scores (STAI-state, STAI-trait, and BDI-II). Correlations were computed using electrodes O1, O2, P7 and P8.

## Discussion

This study aimed to elucidate whether adults with a history of childhood interpersonal trauma show a cortical E/I imbalance, indicated by EEG signal analysis, across resting-state and a visual reaction-time task. Considering existing findings from animal models (35–37, 45) and outcomes of preliminary studies using ^1^HMRS spectroscopy in patients burdened with early adversity (48), we hypothesized that applied computational methods will show significant E/I balance alterations in a non-clinical group of young adults. Additionally, it was assumed that expected task-related alterations will be observable principally in the anterior cortical regions, having in mind previously described findings suggesting various abnormalities in the frontal cortex activity (23). Generally, obtained results corroborated these hypotheses.

In terms of behavioral variables, group comparisons showed significant differences in both affect and task performance measures. The group with intensified traumatic exposure (high-trauma group) scored higher in questionnaires evaluating depressiveness, as well as both trait and anxiety state. Higher levels of these negative emotions are commonly reported findings in this specific group (58, 59). In case of task performance, there were no significant differences in the mean reaction time, nor the *tau* index covering a substantial response time variability typical for the clinical group with attentional disturbances (72). The difference considered a *sigma* parameter that reflects the variability of the normal component with faster and more regular responses. Referring to Leth-Steensen et al. (73) seminal paper on ex-Gaussian markers interpretation, significantly higher *sigma* scores in the high-trauma group may indicate some flaws in response preparation. Apart from it, the response accuracy in the short interval task block was also significantly lower in the high-trauma group. The above presented outcome suggests that participants from the high-trauma group exhibited mild, but detectable impairments in task performance.

Group comparisons of the resting-state EEG revealed significant differences in aperiodic 1/f slope, characterized by a flattening of the power spectrum in the high-trauma group. This effect was primarily observed over posterior electrodes. Additionally, an inverse model predicted a significant increase in the E/I ratio within the same posterior regions for the high-trauma group. Hence, both computational methods clearly indicate a shift toward neuronal increased excitation (and/or reduced inhibition) in the high-trauma sample. We also conducted a direct group comparison of measures derived from task-state EEG data, including the 1/f spectral slope and inferred E/I balance, computed during the pre- and post-stimulus intervals, and observed only a small number of significant, highly localized effects, restricted to the long-interval task block.

Complementing the inter-group results described above, analyses of pre- to post-stimulus E/I balance reconfiguration revealed substantial group effects. According to previous findings in human recordings (57, 74), task-related events like stimulus onset shift the aperiodic activity into steeper values, which is consistent with a global increase in inhibitory activity representing the momentary blockage of ongoing processing, facilitating stimulus-specific processing. This shift is usually broadband and has widespread scalp distribution. Moreover, studies in animal models have shown that inhibitory activity dominates sensory responses in the awake cortex and critically shapes cortical dynamics (75, 76). Comparison of aperiodic 1/f slope and model-derived E/I ratio before (- 500 ms) and after (+500 ms) stimulus onset in the low-trauma group fully confirms the above-described E/I ratio dynamics, i.e., the post-stimulus window is dominated by an inhibitory spectrum almost over the entire cortex. This effect applies to both task blocks. Such reconfiguration in the high-trauma group was only partial and did not involve the anterior cortical regions, especially the prefrontal cortex (PFC). This effect occurred to a similar extent for both task blocks.

Analyses of correlations between resting-state E/I markers and questionnaire-based measures of depressiveness and anxiety revealed no significant effects. Given the pronounced group differences in affective state, the absence of such associations, particularly within the high-trauma group, suggests that the observed between-group differences in the 1/f slope and modeled E/I ratio are unlikely to be driven by current levels of anxiety or depressiveness.

### Implications

To our knowledge, this is the first study to assess neurophysiologically derived neuronal E/I balance in a non-clinical population with a history of early adversity. These findings extend prior work using ^1^H-MRS markers of excitation and inhibition, which focused on localized brain regions involved in emotion regulation, such as the anterior cingulate cortex (48). In contrast, our results demonstrate that group differences in E/I balance are distributed across broad cortical regions and are evident even in case of a simple, non-affective task. Notably, significant effects in earlier ^1^H-MRS studies were often reported in clinically affected samples (46, 47), with trauma–Glx associations frequently driven by the inclusion of clinically depressed individuals (48). Collectively, our findings indicate that E/I imbalance associated with early stress is not confined to resting-state conditions or emotion-related regions and can be detected in a non-clinical group.

The observed shift toward increased excitation and/or reduced inhibition in the high-trauma group suggests that early adversity disrupts core neurophysiological mechanisms and alters normative developmental trajectories. Recent studies indicate that the EEG-based aperiodic slope follows a U-shaped trajectory across age, with a minimum observed around 25–30 years, corresponding to the transition from young adulthood to middle adulthood (77). Given the age range of our sample, the between-group differences observed here are therefore most consistent with trauma-related alterations in functional brain development. Typical development is also characterized by task-related steepening of the aperiodic slope, alongside a gradual dominance of inhibition from childhood to early adulthood, particularly in prefrontal regions (77, 78). Accordingly, evidence from PFC maturation studies shows that improvement of cognitive functioning during adolescence is linked to reduced spontaneous glutamatergic activity and strengthened inhibitory mechanisms (79). Within this framework, the absence of task-related prefrontal inhibitory reactivity in the high-trauma group aligns with extensive evidence of PFC dysfunction following early adversity. A recent meta-analysis identified PFC impairment as the most consistent finding across fMRI studies of adversity, alongside amygdala hyperactivity (15). Reduced task-related inhibition within PFC may therefore lead to elevated spontaneous activity, diminished signal-to-noise ratio, and impaired cognitive control, contributing to the inconsistent performance observed in the high-trauma group.

Early adversity is a well-established risk factor for a wide range of psychiatric outcomes (80, 81), and the present findings suggest that trauma-related E/I imbalance may constitute a latent neurobiological phenotype that increases susceptibility to psychopathology. Persistent excitation dominance at rest, combined with deficient inhibitory modulation during cognitive demands, may potentially contribute to numerous behavioral consequences; hence, in this sense, altered E/I balance may function as an intermediate phenotype linking early environmental stressors to later clinical manifestations. Given the high prevalence of childhood adversity among psychiatric populations (82), neural differences attributed to diagnostic status in case–control studies may partly reflect unmeasured effects of early trauma. Therefore, we suggest that systematic assessment of early adversity should thus be considered essential in studies examining neural markers of neuropsychiatric disorders.

The observed increased E/I balance in trauma-exposed young adults has important clinical and transdiagnostic implications. Although the studied individuals did not meet diagnostic criteria for psychiatric disorders, the identified neurophysiological profile closely resembles patterns reported in some neuropsychiatric conditions, including post-traumatic stress disorder (83, 84), major depressive disorder (85, 86), and generalized epilepsy (87). This convergence suggests that E/I dysregulation may represent a shared vulnerability mechanism rather than a disorder-specific neural alteration.

### Limitations

First, the cross-sectional design precludes causal inferences regarding the relationship between early psychological trauma and alterations in E/I balance. Longitudinal studies are therefore required to determine whether the observed E/I shifts reflect enduring developmental consequences of early adversity or adaptive responses that evolve over time. Second, trauma exposure was assessed retrospectively using a self-report questionnaire, which may be subject to recall bias and limits precision regarding the timing and specific characteristics of adverse experiences. Current perspectives on trauma–brain relationships suggest that neurobiological consequences may depend on early adversity type, such as whether adversity primarily reflects threat or neglect (88). Such typological differentiation could not be examined in the present study due to the structure of the questionnaire, which yields a composite score encompassing multiple forms of adversity. To model a neurophysiological proxy of E/I balance, we applied two partially independent methods to strengthen the robustness of the interpretations. Nevertheless, regarding the aperiodic slope, although this parameter has been largely validated (79, 89), some studies examining its relationship with direct measures of GABAergic and glutamatergic function have failed to confirm the expected associations (90). In addition, reliance on the aperiodic 1/f slope alone may conflate distinct neural mechanisms, underscoring the need for multimodal or model-constrained approaches when interpreting E/I balance (52). Some limitations of the modeling approach should also be acknowledged. The current framework relies on a local model of cortical dynamics and does not explicitly incorporate cortico–cortical connectivity, despite substantial evidence that long-range interactions play a critical role in shaping certain neural processes (e.g., low-frequency neuronal oscillations). Future work should therefore extend this modeling scheme to incorporate large-scale network connectivity and its potential influence on early trauma–related neural dynamics.

## Conclusions

The present study provides converging evidence that exposure to ChT is associated with persistent alterations in neurophysiological cortical E/I balance, manifested as a shift toward excitation at rest and incomplete task-related inhibitory modulation, particularly in frontal cortical regions. These neurophysiological alterations were observed in a non-clinical, drug-naïve sample and were not accounted for by current levels of anxiety or depressiveness, suggesting that they reflect trait-like consequences of early adversity rather than transient affective states. The similarity between the observed E/I profiles and those reported in several psychiatric conditions supports the notion that altered E/I balance may represent a transdiagnostic vulnerability marker shaped by early-life stress.

## References

1. F. Devi et al., The prevalence of childhood trauma in psychiatric outpatients. Ann Gen Psychiatry 18, 15 (2019).

2. C. P. Carr, C. M. Martins, A. M. Stingel, V. B. Lemgruber, M. F. Juruena, The role of early life stress in adult psychiatric disorders: a systematic review according to childhood trauma subtypes. J Nerv Ment Dis 201, 1007–1020 (2013).

3. E. McCrory, S. A. De Brito, E. Viding, Research review: the neurobiology and genetics of maltreatment and adversity. J Child Psychol Psychiatry 51, 1079–1095 (2010).

4. D. Cross, N. Fani, A. Powers, B. Bradley, Neurobiological Development in the Context of Childhood Trauma. Clin Psychol (New York) 24, 111–124 (2017).

5. K. R. Kuhlman, I. Vargas, E. G. Geiss, N. L. Lopez-Duran, Age of Trauma Onset and HPA Axis Dysregulation Among Trauma-Exposed Youth. J Trauma Stress 28, 572–579 (2015).

6. Y. Dvir, J. D. Ford, M. Hill, J. A. Frazier, Childhood maltreatment, emotional dysregulation, and psychiatric comorbidities. Harv Rev Psychiatry 22, 149–161 (2014).

7. A. Dodaj, M. Krajina, K. Sesar, N. Šimić, The Effects of Maltreatment in Childhood on Working Memory Capacity in Adulthood. Eur J Psychol 13, 618–632 (2017).

8. M. Majer, U. M. Nater, J. M. Lin, L. Capuron, W. C. Reeves, Association of childhood trauma with cognitive function in healthy adults: a pilot study. BMC Neurol 10, 61 (2010).

9. A. J. Petkus, E. J. Lenze, M. A. Butters, E. W. Twamley, J. L. Wetherell, Childhood Trauma Is Associated With Poorer Cognitive Performance in Older Adults. J Clin Psychiatry 79 (2018).

10. H. B. Daníelsdóttir et al., Adverse Childhood Experiences and Adult Mental Health Outcomes. JAMA Psychiatry 81, 586–594 (2024).

11. A. Bhattarai et al., Childhood Adversity and Mental Health Outcomes Among University Students: A Longitudinal Study. Can J Psychiatry 68, 510–520 (2023).

12. S. Salzmann, M. Salzmann-Djufri, F. Euteneuer, Childhood Emotional Neglect and Cardiovascular Disease: A Narrative Review. Front Cardiovasc Med 9, 815508 (2022).

13. M. J. H. Begemann et al., Childhood trauma is associated with reduced frontal gray matter volume: a large transdiagnostic structural MRI study. Psychol Med 53, 741–749 (2023).

14. R. A. Madden et al., Structural brain correlates of childhood trauma with replication across two large, independent community-based samples. Eur Psychiatry 66, e19 (2023).

15. N. Hosseini-Kamkar et al., Adverse Life Experiences and Brain Function: A Meta-Analysis of Functional Magnetic Resonance Imaging Findings. JAMA Netw Open 6, e2340018 (2023).

16. Y. Hakamata, Y. Suzuki, H. Kobashikawa, H. Hori, Neurobiology of early life adversity: A systematic review of meta-analyses towards an integrative account of its neurobiological trajectories to mental disorders. Front Neuroendocrinol 65, 100994 (2022).

17. N. S. Philip et al., Exposure to childhood trauma is associated with altered n-back activation and performance in healthy adults: implications for a commonly used working memory task. Brain Imaging Behav 10, 124–135 (2016).

18. C. H. Allen et al., Aberrant resting-state functional connectivity associated with childhood trauma among juvenile offenders. Neuroimage Clin 37, 103343 (2023).

19. M. D. De Bellis, A. Zisk, The biological effects of childhood trauma. Child Adolesc Psychiatr Clin N Am 23, 185–222, vii (2014).

20. S. H. Lee, Y. Park, M. J. Jin, Y. J. Lee, S. W. Hahn, Childhood Trauma Associated with Enhanced High Frequency Band Powers and Induced Subjective Inattention of Adults. Front Behav Neurosci 11, 148 (2017).

21. F. M. Howells, D. J. Stein, V. A. Russell, Childhood Trauma is Associated with Altered Cortical Arousal: Insights from an EEG Study. Front Integr Neurosci 6, 120 (2012).

22. G. Zukerman, L. Fostick, E. Ben-Itzchak, Early automatic hyperarousal in response to neutral novel auditory stimuli among trauma-exposed individuals with and without PTSD: An ERP study. Psychophysiology 55, e13217 (2018).

23. R. Kessler et al., Long-Term Neuroanatomical Consequences of Childhood Maltreatment: Reduced Amygdala Inhibition by Medial Prefrontal Cortex. Front Syst Neurosci 14, 28 (2020).

24. S. Kim, J. S. Kim, M. J. Jin, C. H. Im, S. H. Lee, Dysfunctional frontal lobe activity during inhibitory tasks in individuals with childhood trauma: An event-related potential study. Neuroimage Clin 17, 935–942 (2018).

25. S. Zhang et al., In vivo whole-cortex marker of excitation-inhibition ratio indexes cortical maturation and cognitive ability in youth. Proc Natl Acad Sci U S A 121, e2318641121 (2024).

26. A. L. Dorrn, K. Yuan, A. J. Barker, C. E. Schreiner, R. C. Froemke, Developmental sensory experience balances cortical excitation and inhibition. Nature 465, 932–936 (2010).

27. V. O. Manyukhina et al., Globally elevated excitation-inhibition ratio in children with autism spectrum disorder and below-average intelligence. Mol Autism 13, 20 (2022).

28. S. Trakoshis et al., Intrinsic excitation-inhibition imbalance affects medial prefrontal cortex differently in autistic men versus women. Elife 9, e55684 (2020).

29. H. Bruining et al., Measurement of excitation-inhibition ratio in autism spectrum disorder using critical brain dynamics. Sci Rep 10, 9195 (2020).

30. M. Chini, T. Pfeffer, I. Hanganu-Opatz, An increase of inhibition drives the developmental decorrelation of neural activity. Elife 11 (2022).

31. M. Chini, I. L. Hanganu-Opatz, Prefrontal Cortex Development in Health and Disease: Lessons from Rodents and Humans. Trends Neurosci 44, 227–240 (2021).

32. B. Larsen, B. Luna, Adolescence as a neurobiological critical period for the development of higher-order cognition. Neurosci Biobehav Rev 94, 179–195 (2018).

33. A. Saberi, et al., Adolescent maturation of cortical excitation-inhibition balance based on individualized biophysical network modeling. *bioRxiv* (2024).

34. S. L. Shamansky, G. H. Glaser, Socioeconomic characteristics of childhood seizure disorders in the New Haven area: an epidemiologic study. Epilepsia 20, 457–474 (1979).

35. L. E. Wearick-Silva, A. D. Sebben, Z. S. M. Costa-Ferro, D. R. Marinowic, M. L. Nunes, Undernourishment and recurrent seizures early in life impair Long-Term Potentiation and alter NMDAR and AMPAR expression in rat hippocampus. Int J Dev Neurosci 75, 13–18 (2019).

36. G. Kumar et al., Early life stress enhancement of limbic epileptogenesis in adult rats: mechanistic insights. PLoS One 6, e24033 (2011).

37. L. Velísek, K. Jehle, S. Asche, J. Velísková, Model of infantile spasms induced by N-methyl-D-aspartic acid in prenatally impaired brain. Ann Neurol 61, 109–119 (2007).

38. C. M. Dubé et al., Hyper-excitability and epilepsy generated by chronic early-life stress. Neurobiol Stress 2, 10–19 (2015).

39. A. Hooper, R. Paracha, J. Maguire, Seizure-induced activation of the HPA axis increases seizure frequency and comorbid depression-like behaviors. Epilepsy Behav 78, 124–133 (2018).

40. K. K. O’Toole, A. Hooper, S. Wakefield, J. Maguire, Seizure-induced disinhibition of the HPA axis increases seizure susceptibility. Epilepsy Res 108, 29–43 (2014).

41. X. Bian, W. Yang, J. Lin, B. Jiang, X. Shao, Hypothalamic-Pituitary-Adrenal Axis and Epilepsy. J Clin Neurol 20, 131–139 (2024).

42. A. C. Wulsin, M. B. Solomon, M. D. Privitera, S. C. Danzer, J. P. Herman, Hypothalamic-pituitary-adrenocortical axis dysfunction in epilepsy. Physiol Behav 166, 22–31 (2016).

43. L. Jacobson, R. Sapolsky, The role of the hippocampus in feedback regulation of the hypothalamic-pituitary-adrenocortical axis. Endocr Rev 12, 118–134 (1991).

44. J. P. Herman et al., Evidence for hippocampal regulation of neuroendocrine neurons of the hypothalamo-pituitary-adrenocortical axis. J Neurosci 9, 3072–3082 (1989).

45. H. Karst et al., Acceleration of GABA-switch after early life stress changes mouse prefrontal glutamatergic transmission. Neuropharmacology 234, 109543 (2023).

46. L. A. Averill et al., Early life stress and glutamate neurotransmission in major depressive disorder. Eur Neuropsychopharmacol 35, 71–80 (2020).

47. A. I. Sonmez, et al., A pilot spectroscopy study of adversity in adolescents. Biomark Neuropsychiatry 5 (2021).

48. L. Herrmann et al., Cross-sectional study of retrospective self-reported childhood emotional neglect and inhibitory neurometabolite levels in the pregenual anterior cingulate cortex in adult humans. Neurobiol Stress 25, 100556 (2023).

49. P. Hepsomali et al., Signatures of exposure to childhood trauma in young adults in the structure and neurochemistry of the superior temporal gyrus. J Psychopharmacol 37, 510–519 (2023).

50. L. M. Levy, A. J. Degnan, GABA-based evaluation of neurologic conditions: MR spectroscopy. AJNR Am J Neuroradiol 34, 259–265 (2013).

51. J. Ahmad et al., From mechanisms to markers: novel noninvasive EEG proxy markers of the neural excitation and inhibition system in humans. Transl Psychiatry 12, 467 (2022).

52. A. Orozco Valero et al., A Python toolbox for neural circuit parameter inference. NPJ Syst Biol Appl 11, 45 (2025).

53. A. Ziaeemehr et al., Virtual Brain Inference (VBI), a flexible and integrative toolkit for efficient probabilistic inference on whole-brain models. Elife 14 (2025).

54. P. Martínez-Cañada et al., Combining aperiodic 1/f slopes and brain simulation: An EEG/MEG proxy marker of excitation/inhibition imbalance in Alzheimer’s disease. Alzheimer’s & Dementia: Diagnosis, Assessment & Disease Monitoring 15, e12477 (2023).

55. J. L. Molina et al., Memantine Effects on Electroencephalographic Measures of Putative Excitatory/Inhibitory Balance in Schizophrenia. Biol Psychiatry Cogn Neurosci Neuroimaging 5, 562–568 (2020).

56. M. M. Robertson et al., EEG power spectral slope differs by ADHD status and stimulant medication exposure in early childhood. J Neurophysiol 122, 2427–2437 (2019).

57. M. Gyurkovics, G. M. Clements, K. A. Low, M. Fabiani, G. Gratton, Stimulus-induced changes in 1/f-like background activity in EEG. J Neurosci 42, 7144–7151 (2022).

58. D. A. Chu, L. M. Williams, A. W. Harris, R. A. Bryant, J. M. Gatt, Early life trauma predicts self-reported levels of depressive and anxiety symptoms in nonclinical community adults: relative contributions of early life stressor types and adult trauma exposure. J Psychiatr Res 47, 23–32 (2013).

59. J. G. Hovens et al., Impact of childhood life events and trauma on the course of depressive and anxiety disorders. Acta Psychiatr Scand 126, 198–207 (2012).

60. M. Deodato, D. Melcher, P. Vuilleumier, Fearful Events Induce Sustained Changes in Spontaneous Aperiodic EEG. Psychophysiology 62, e70103 (2025).

61. M. Z. Gehred et al., Long-term Neural Embedding of Childhood Adversity in a Population-Representative Birth Cohort Followed for 5 Decades. Biol Psychiatry 90, 182–193 (2021).

62. E. Hagen et al., Brain signal predictions from multi-scale networks using a linearized framework. PLoS Comput Biol 18, e1010353 (2022).

63. Z. Juczyński, N. Ogińska-Bulik, Measurement of post-traumatic stress disorder: Polish version of the Impact of Event Scale–Revised. Psychiatria 6, 10 (2009).

64. D. P. Bernstein, L. Fink, Childhood Trauma Questionnaire: A retrospective self-report questionnaire and manual (The Psychological Corporation, San Antonio, TX, 1998).

65. A. T. Beck, R. A. Steer, G. K. Brown, Manual for the Beck Depression Inventory–II (The Psychological Corporation, San Antonio, TX, 1996).

66. C. D. Spielberger, Manual for the State-Trait Anxiety Inventory (STAI) (Consulting Psychologists Press, Palo Alto, CA, 1983).

67. P. R. Killeen, Absent without leave; a neuroenergetic theory of mind wandering. Front Psychol 4, 373 (2013).

68. F. C. Fortenbaugh et al., Interpersonal early-life trauma alters amygdala connectivity and sustained attention performance. Brain Behav 7, e00684 (2017).

69. T. Donoghue et al., Parameterizing neural power spectra into periodic and aperiodic components. Nat Neurosci 23, 1655–1665 (2020).

70. T. Donoghue, A Systematic Review of Aperiodic Neural Activity in Clinical Investigations. Eur J Neurosci 62, e70255 (2025).

71. Y. Lacouture, D. Cousineau, How to use MATLAB to fit the ex-Gaussian and other probability functions to a distribution of response times. Tutorials in Quantitative Methods for Psychology 4, 10 (2008).

72. M. Bella-Fernández, M. Martin-Moratinos, C. Li, P. Wang, H. Blasco-Fontecilla, Differences in Ex-Gaussian Parameters from Response Time Distributions Between Individuals with and Without Attention Deficit/Hyperactivity Disorder: A Meta-analysis. Neuropsychol Rev 34, 320–337 (2024).

73. C. Leth-Steensen, Z. King Elbaz, V. I. Douglas, Mean response times, variability, and skew in the responding of ADHD children: a response time distributional approach. Acta Psychologica 104, 167–190 (2000).

74. P. Kałamała et al., Event-induced modulation of aperiodic background EEG: Attention-dependent and age-related shifts in E:I balance, and their consequences for behavior. Imaging Neurosci (Camb) 2 (2024).

75. J. Veit, R. Hakim, M. P. Jadi, T. J. Sejnowski, H. Adesnik, Cortical gamma band synchronization through somatostatin interneurons. Nat Neurosci 20, 951–959 (2017).

76. B. Haider, M. Häusser, M. Carandini, Inhibition dominates sensory responses in the awake cortex. Nature 493, 97–100 (2013).

77. Z. R. Cross et al., The development of aperiodic neural activity in the human brain. Nat Hum Behav 9, 2548–2563 (2025).

78. A. Saberi et al., Adolescent maturation of cortical excitation-inhibition ratio based on individualized biophysical network modeling. Sci Adv 11, eadr8164 (2025).

79. S. D. McKeon et al., Aperiodic EEG and 7T MRSI evidence for maturation of E/I balance supporting the development of working memory through adolescence. Dev Cogn Neurosci 66, 101373 (2024).

80. E. Giampetruzzi et al., Impact of adverse childhood experiences on risk for internalizing psychiatric disorders in youth at clinical high-risk for psychosis. Psychiatry Res 342, 116214 (2024).

81. Y. Kim, K. Kim, K. G. Chartier, T. L. Wike, S. E. McDonald, Adverse childhood experience patterns, major depressive disorder, and substance use disorder in older adults. Aging Ment Health 25, 484–491 (2021).

82. R. Knipschild et al., Childhood adversity in a youth psychiatric population: prevalence and associated mental health problems. Eur J Psychotraumatol 15, 2330880 (2024).

83. N. Kovacevic, A. Meghdadi, C. Berka, Characterizing PTSD Using Electrophysiology: Towards A Precision Medicine Approach. Clin EEG Neurosci 56, 305–315 (2025).

84. M. P. Deiber et al., A biomarker of brain arousal mediates the intergenerational link between maternal and child post-traumatic stress disorder. J Psychiatr Res 177, 305–313 (2024).

85. S. E. Woronko et al., Alterations in aperiodic neural activity associated with major depressive disorder. Nature Mental Health 3, 1181–1190 (2025).

86. A. Zandbagleh, S. Sanei, H. Azami, Implications of Aperiodic and Periodic EEG Components in Classification of Major Depressive Disorder from Source and Electrode Perspectives. Sensors (Basel) 24 (2024).

87. M. Kopf et al., Aperiodic Activity Indexes Neural Hyperexcitability in Generalized Epilepsy. eNeuro 11 (2024).

88. K. A. McLaughlin, M. A. Sheridan, H. K. Lambert, Childhood adversity and neural development: deprivation and threat as distinct dimensions of early experience. Neurosci Biobehav Rev 47, 578–591 (2014).

89. A. Sheldon et al., Aperiodic EEG activity correlates with occipital glutamate from 7 tesla MRS. Journal of Vision 25, 2151 (2025).

90. S. V. Salvatore et al., Periodic and aperiodic changes to cortical EEG in response to pharmacological manipulation. J Neurophysiol 131, 529–540 (2024).

