## Supplementary figures and images for "Neurophysiological excitation/inhibition imbalance in young adults burdened with childhood interpersonal trauma"

### Supplemental Figure 1

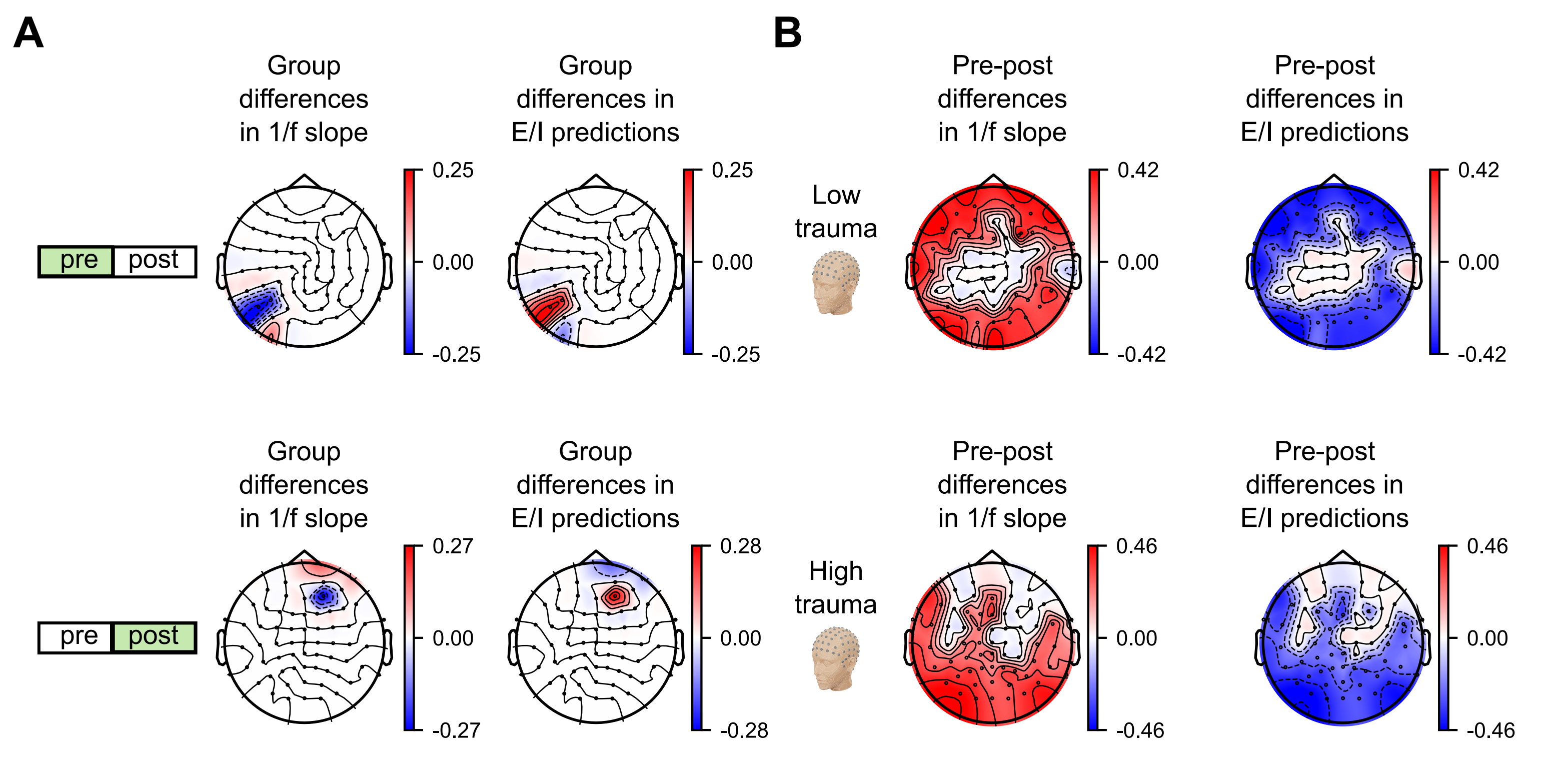
